# Single-cell Landscape of T Cell Heterogeneity in Kawasaki Disease: STAT3/JAK Axis Regulates the Lineage Differentiation Bias of Th17 Cells

**DOI:** 10.64898/2026.03.18.712795

**Authors:** Sirui Song, Yanfang Zong, Yanbing Xu, Liqin Chen, Yuanyuan Zhou, Le Chen, Guang Li, Tingting Xiao, Min Huang

## Abstract

**Background:** Kawasaki disease (KD) is a pediatric systemic vasculitis in which T-cell-mediated immune responses play a pivotal role. However, the precise dynamic evolution of T-cell subsets during disease progression remains poorly understood.

**Methods:** Single-cell RNA sequencing (scRNA-seq) was employed to perform high-resolution annotation of peripheral blood mononuclear cells (PBMCs) from healthy controls and KD patients, both pre- and post- IVIG treatment. T-cell developmental trajectories were reconstructed via Monocle3-based pseudotime analysis. Furthermore, the functional significance of the significant pathway was validated in a CAWS-induced KD murine model.

**Results:** A high-resolution single-cell landscape identified 13 distinct T-cell subtypes. Pseudotime analysis revealed a significant lineage commitment of CD4^+^ T cells toward a Th17 phenotype during the acute phase of KD, synchronized with the transcriptional upregulation of the STAT3/JAK signaling axis. Animal experiments further demonstrated that pharmacological inhibition of this pathway substantially attenuated inflammatory infiltration in the cardiac vasculature of KD mice.

**Conclusion:** This study identifies the STAT3/JAK-mediated Th17 differentiation bias as a potential regulatory program associated with acute inflammation in Kawasaki disease, thereby highlighting the STAT3/JAK axis as a potential therapeutic target.

## 1. Background

Kawasaki Disease (KD) is a pediatric systemic vasculitis characterized by widespread inflammation of the blood vessels. It remains a leading cause of acquired heart disease in children, posing a significant threat to pediatric health. Since its initial report in 1967, the global incidence of KD has risen annually, particularly within developed nations where it has become the primary etiology of childhood acquired heart disease ^[1]^. Clinical manifestations predominantly include fever, mucocutaneous lesions, and lymphadenopathy. If left untreated, KD may lead to severe coronary artery lesions (CALs), which can be life-threatening ^[2]^. Although the precise etiology remains elusive, current evidence suggests that the pathogenesis involves an overactivation of the immune system triggered by a combination of infectious agents and genetic susceptibility. Currently, the standard treatment regimen consists of intravenous immunoglobulin (IVIG) combined with aspirin; however, 10% to 20% of patients exhibit a lack of response to initial therapy, termed IVIG resistance ^[3]^. Consequently, exploring the immunological mechanisms—specifically the role of T cells—is critical for elucidating the disease etiology and optimizing therapeutic strategies.

T cells play a pivotal role in the immunopathogenesis of KD. Previous studies have demonstrated significant shifts in the distribution and function of T-cell subsets, which correlate closely with disease severity and treatment response ^[4]^. For instance, a notable increase in Th17 cells within the CD4^+^ T-cell population has been observed during the acute phase of KD, alongside a relative reduction in regulatory T cells (Tregs). This Th17/Treg cells imbalance is considered a key factor driving persistent inflammation and IVIG resistance ^[5]^. Furthermore, CD8^+^ T cells contribute significantly to vascular inflammation in KD, where their overactivation can lead to endothelial damage and exacerbated inflammatory responses ^[6]^. Despite existing research characterizing T-cell subsets in the peripheral blood and lesion tissues of KD patients, a systematic understanding of their dynamic transitions and functional variations across different stages—acute, subacute, and convalescent—remains insufficient ^[7]^. Recent advancements in single-cell RNA sequencing (scRNA-seq) have revolutionized immunology by providing unparalleled resolution for dissecting cellular heterogeneity. Previous efforts by this research team established an initial immune cell atlas of acute KD, analyzing cell-cell communication and specific expression modules, which provided a foundation for understanding acute-phase immune dynamics ^[8–10]^. Nevertheless, a gap remains in the systematic annotation of T-cell subsets in KD, particularly regarding their longitudinal evolution and functional shifts. The present study utilizes scRNA-seq to perform a comprehensive annotation of T-cell subsets in KD patients, revealing their temporal dynamics and gene expression profiles. These findings are further validated through a murine model treated with Th17 inhibitors to confirm the functional significance of the identified pathways.

## 2. Methods

### 2.1 Study Population

Pediatric patients diagnosed with KD and admitted to the Department of cardiology,Shanghai Children’s Hospital, School of Medicine, Shanghai Jiao Tong University between December 2019 and December 2021 were enrolled in this study. The diagnostic criteria for the KD group followed the 2017 American Heart Association (AHA) guidelines ^[11]^, all patients tested negative for SARS-CoV-2. Patients presenting with fewer than four features were classified as having incomplete KD. Upon diagnosis, patients received high-dose intravenous immunoglobulin (IVIG, 1 g/kg/day for 2 consecutive days) in combination with oral aspirin (30 mg/kg/day). Blood samples were collected from all KD patients at consistent time points: the pre-treatment (KD-Before) samples were obtained on the day of diagnosis prior to IVIG administration, and the post-treatment (KD-After) samples were collected 24 hours after the stabilization of body temperature following IVIG therapy.The healthy control (HC) group consisted of age-matched children (under 6 years old) recruited from the outpatient physical examination center. Inclusion criteria for the HC group required no recent history of fever, infection, or vaccination. Exclusion criteria for both groups included a prior history of KD or any pre-existing autoimmune diseases.This study was reviewed and approved by the Ethics Committee of the Shanghai Children’s Hospital. Informed consent was obtained from the legal guardians of all participating children.

### 2.2 Single-Cell Preparation and Sequencing

Venous blood (2 mL) was collected from each donor using EDTA-anticoagulated tubes. Fresh blood samples were immediately processed to prepare peripheral blood mononuclear cell (PBMC) suspensions. The prepared cell suspensions were required to meet the following quality control criteria: cell viability > 85%, total cell count > 200,000, and a cell diameter range of 7–60 um. Additionally, the suspensions had to be free of erythrocytes, cell aggregates, visible debris, and clumps.Single-cell library construction was initiated within 30 minutes of PBMC suspension preparation. Using the 10x Genomics Chromium platform (5’ library construction method), both 5’ gene expression (GEX) and V(D)J sequence libraries were established. Sequencing was performed on the Illumina NovaSeq platform in PE150 mode. The target cell recovery for each sample ranged from 1,000 to 10,000 cells. The sequencing depth was set at approximately 30M reads/cell for 5’ gene expression and 3M reads/cell for V(D)J sequencing.

### 2.3 Bioinformatics Analysis

#### 2.3.1 Data Preprocessing, Quality Control, and Preliminary Annotation

The raw single-cell RNA sequencing (scRNA-seq) data were processed using the Seurat (v5.3.1) R package ^[12, 13]^. Quality control (QC) was performed to exclude low-quality cells based on standard metrics: cells with fewer than 200 or more than 5,000 detected genes (nFeature_RNA), or a mitochondrial gene percentage (percent.mt) exceeding 10% were removed. The resulting count matrices were normalized using the LogNormalize method with a scale factor of 10,000. Subsequently, 2,000 Highly Variable Genes (HVGs) were identified via variance-stabilizing transformation (vst method) for downstream analysis.

To reduce data dimensionality, all genes were scaled and centered (ScaleData), followed by Principal Component Analysis (PCA) performed based on the HVGs. Potential doublets were identified and removed using the DoubletFinder(v2.0.6) R package^[14]^. To account for variations across different batches, the Harmony algorithm was applied to integrate the PCA space and correct for batch effects^[15]^. Cell clustering was performed using the Louvain community detection algorithm based on a Shared Nearest Neighbor (SNN) graph constructed in the PCA space. The Euclidean distance was employed as the metric for neighbor search.. The stability of clusters at various resolutions was evaluated using the clustree(v0.5.1) package to determine the optimal resolution for cell partitioning^[16]^.

Finally, major immune cell populations—including “T/NK cells”, “B cells”, “Plasma cells”, “Neutrophils”, “Monocytes”, “Dendritic Cells (DCs)”, “Platelets”, and “Erythroblasts“—were annotated by examining the expression of canonical lineage markers across each cluster. Specifically, to achieve a high-resolution classification of T-cell subsets, a cluster containing a mixture of T and NK cells (T_NK_mixed) was subsetted and subjected to a subsequent refined annotation pipeline.

#### 2.3.2 Hierarchical and Refined Annotation of T-cell Subsets

To precisely dissect T-cell heterogeneity, a hierarchical cell annotation strategy was implemented. Initially, all T cells were extracted from the preliminary annotation results to form an independent subset. This subset was then subjected to a new round of normalization, dimensionality reduction, batch correction, and clustering analysis.The clustering resolution was optimized to ensure that all retained clusters were characterized by distinct marker gene profiles, thereby ensuring the stability and biological relevance of the downstream analysis.

##### (1) Partitioning of CD4^+^, CD8^+^, and Other T-cell Lineages

T cells were partitioned into CD4^+^, CD8^+^, and other T-cell lineages (primarily γδ T cells) based on cluster-level expression of canonical markers. To resolve T-cell heterogeneity, high-resolution sub-clustering was performed on each lineage. Sub-populations were annotated using established transcriptomic signatures for naive, memory, and effector states, as well as lineage-defining transcription factors and cytokine receptors for helper T and regulatory T subsets. This hierarchical strategy ensured that each cluster was identified by its global transcriptional profile rather than single-gene thresholds.

##### (2) Annotation of Refined T-cell Subsets

The previously extracted CD4^+^ T-cell sub-populations were again subjected to independent high-dimensional reduction and unsupervised clustering. The annotation of resulting clusters was based on combinations of established canonical marker genes. For clusters exhibiting high heterogeneity—specifically mixed populations of helper T cells (Th) and effector memory T cells (TEM)—a strategy of “subset extraction followed by re-clustering” was employed. This involved merging relevant clusters and conducting a third round of independent analysis to achieve refined resolution of these complex subsets. Similarly, CD8^+^ T-cell and CD4^−^CD8^−^ (double-negative) T-cell subsets underwent independent rounds of dimensionality reduction and clustering. γδ T (gDT) cells were further subdivided into naive and effector subpopulations based on their differentiation status.

#### 2.3.3 Cell-Cell Communication Analysis

To systematically infer and characterize the intercellular interaction networks across experimental groups, the CellChat (v2.2.0) R package was employed ^[17]^. Analysis was performed independently for each group to delineate group-specific communication landscapes.

For each group, Seurat objects—following the removal of “mixed” cell populations(e.g., potential doublets and low-quality cells characterized by the absence of distinct lineage markers)—were converted into CellChat objects. Ligand-receptor interactions were predicted based on the CellChatDB.human database. Signaling molecule expression data were extracted and screened for overexpressed genes and interactions. To enhance signal robustness and compensate for technical sparsity in scRNA-seq data, the expression profiles were smoothed by integrating the human protein-protein interaction (PPI) network. The core communication probabilities were inferred using a statistical model based on the law of mass action. To minimize the influence of outliers, a 5% truncated mean was applied to the expression levels within each cell cluster (via the trim parameter). Furthermore, the interaction strengths were adjusted to account for the relative proportion of each cell subpopulation (population.size = TRUE). The resulting networks were subsequently filtered to retain interactions where ligands or receptors were expressed in at least 10 cells.

Finally, communication probabilities were aggregated from individual ligand-receptor pairs to the signaling pathway level. A global communication network was then constructed by consolidating all relevant interactions. To identify key regulatory nodes, network centrality metrics were computed, allowing for the quantitative assessment of each cell subpopulation’s role as a prominent signal sender or receiver.

#### 2.3.4 Pseudotime Trajectory and Cellular Dynamics Analysis

##### (1) Simulation of Cellular Pseudotime States

To investigate developmental and differentiation dynamics within T-cell subsets, pseudotime trajectory analysis was performed with Monocle3 (v1.4.26) ^[18]^ for each lineage (CD4+, CD8+, γδ T cells) and each group. Seurat-derived cells were imported to create CellDataSet objects, re-clustered with the Leiden algorithm, and ordered along a principal graph learned in UMAP space. Naive T-cell subsets were designated as the root, and pseudotime values were computed for all cells. Trajectory-dependent dynamic genes(TDDGs) were identified with the graph_test spatial correlation test and compared across conditions.

##### (2) Co-expression Module Analysis

TDDGs were partitioned into functional modules using the find_gene_modules function in Monocle3. This process utilized the Louvain community detection algorithm (resolution = 0.01) to cluster genes based on their coordinates in the UMAP-embedded space. The optimal number of modules was determined using the elbow method in combination with biological interpretability to ensure that the identified modules captured distinct transcriptional kinetics.

##### (3) Comparative Analysis of Pseudotime Distribution Dynamics

Pseudotime values for each trajectory were normalized to 0–1, divided into 30 bins, and the relative density of each T-cell subtype within these bins was calculated to compare developmental distributions across groups.

To account for potential pseudoreplication, donor-level validation was performed by aggregating counts per donor (pseudobulk).DESeq (v1.60.0) was employed to estimate dispersion and calculate differential expression significance, ensuring the statistical robustness across individuals of TDDGs.

#### 2.3.5 Differential Gene Expression (DGE) Analysis

Differential gene expression analysis was performed on isolated CD4^+^ T-cell subsets with Seurat. Genes were considered differentially expressed if log2 fold change ≥ 0.25, detected in ≥10 % of cluster cells, and adjusted p-value < 0.05 (Wilcoxon rank-sum test, BH correction).

#### 2.3.6 Functional Enrichment Analysis

GO and KEGG enrichment analyses were carried out with clusterProfiler (v4.10.1) to identify biological processes and pathways associated with the identified gene sets^[19]^.

### 2.4 Animal Model Construction and Protein Expression Assay

#### 2.4.1 Preparation of CAWS

Candida albicans water-soluble fraction (CAWS) was prepared following established protocols ^[20]^. The Candida albicans strain (NBRC: 1385) was purchased from the Institute for Biology in Osaka, Japan. To initiate the preparation, C. albicans was dissolved in sterile water and inoculated into a broth medium (Millipore, Cat# 69966-500G). The culture was incubated at 27°C with continuous shaking at 1,600 rpm overnight. Subsequently, 1 mL of the culture was transferred into 1 L of sterile nutrient solution and incubated at 27°C with shaking at 1,600 rpm for 24 hours.

Following incubation, an equal volume of absolute ethanol was added, and the solution was incubated at 4°C overnight. The mixture was then subjected to centrifugation at 9,000 rpm for 15 minutes at 4°C. The precipitate was collected and rinsed with double-distilled water (ddH2O). The supernatant was retained, mixed with an equal volume of absolute ethanol, and again incubated at 4°C overnight. After a second round of centrifugation at 9,000 rpm for 15 minutes at 4°C, the resulting pellet was lyophilized and weighed. Finally, the precipitate was dissolved in phosphate-buffered saline (PBS) to reach a concentration of 20 mg/mL.

#### 2.4.2 Animal Model Establishment and Intervention

Specific-pathogen-free (SPF) male C57BL/6 mice (n=30, 4 weeks old) were purchased from Shanghai Shengchang Biological Technology Co., Ltd. (Shanghai, China). Ammonium pyrrolidinedithiocarbamate (PDTC) was utilized to block Th17 differentiation ^[21]^. All mice were housed in standard laboratory cages under controlled environmental conditions with a constant temperature (25 ± 2 °C) and humidity (50 % ± 5 %).The C57BL/6 mice were randomly assigned into three groups: the Normal Control group (n=10), the CAWS group (n=10), and the CAWS+PDTC group (n=10). The intervention protocols were administered as follows:Normal Control group: Mice received daily intraperitoneal (i.p.) injections of PBS for 5 consecutive days, followed by daily oral gavage of physiological saline for 28 days.CAWS group: Mice received daily i.p. injections of CAWS (4 mg/day) for 5 consecutive days, followed by daily oral gavage of physiological saline for 28 days.CAWS+PDTC group: Mice received daily i.p. injections of CAWS (4 mg/day) for 5 consecutive days, followed by daily oral gavage of PDTC (20 mg/kg/day; Yuanmu, S30342) for 28 days.All animal procedures were performed in strict accordance with the Guide for the Care and Use of Laboratory Animals published by the National Institutes of Health (NIH).

#### 2.4.3 Preparation of Murine Cardiac Specimens

At the end of the 28-day intervention period, all mice were euthanized via CO2 asphyxiation. For each group, cardiac tissues from three mice were harvested, fixed in 4 % paraformaldehyde, and embedded in paraffin to prepare transverse sections for immunohistochemical staining. The remaining cardiac tissues from each group were immediately snap-frozen in liquid nitrogen and stored at −80 °C for subsequent quantification of IL-17A, SOCS3, RORγt, JAK3, and NF-κB.

#### 2.4.4 Western Blot Analysis

Cardiac levels of IL-17A, STAT3, SOCS3, RORγt, JAK3, and NF-κB were determined by Western blot. Total protein was extracted from six hearts per group using ice-cold lysis buffer (50 mmol/L Tris-Cl pH 7.4, 150 mmol/L NaCl, 1 mmol/L EDTA, 1 % NP40, 0.25 % sodium deoxycholate, 1 mmol/L PMSF, plus protease and phosphatase inhibitors). Sixty-microgram samples were resolved on 10 % SDS-PAGE, transferred to PVDF membranes, blocked, and incubated overnight at 4 °C with primary antibodies against STAT3 (Abcam, ab68153, 1:1000), SOCS3 (Abcam, ab280884, 1:1000), RORγt (Invitrogen, 14-6981-82, 1:1000), JAK3 (Proteintech, 80331-1-RR, 1:1000), NF-κB (CST, 8242S, 1:2000), and IL-17A (Abcam, ab79056, 1:500), using α-tubulin (CST,2125S, 1:2000) or GAPDH (Epizyme Biotech,LF206, 1:1000) as loading controls. Membranes were then incubated with HRP-conjugated goat anti-mouse or anti-rabbit secondary antibodies (Proteintech,HY-P8004;SA00001-2, 1:2000). Bands were visualized with an Amersham Imager 600 and quantified using Image-Pro Plus software.Data are presented as mean ± standard error of the mean (SEM). Statistical analyses were conducted using GraphPad Prism 9.0, and differences among multiple groups were evaluated via one-way analysis of variance (ANOVA). A value of P < 0.05 was considered statistically significant.

## 3. Results

### 3.1 Participant Characteristics

The clinical characteristics of all pediatric participants included in this study are summarized in Table 1. A total of seven patients with KD were enrolled for single-cell RNA sequencing (scRNA-seq). This cohort comprised four males and three females, with a mean age of 3.0 ± 1.4 years (range: 1.6–5.3 years). All patients responded effectively to IVIG therapy and did not develop coronary artery lesions (CALs).

**Table 1.** The demographic and clinical characteristics of children in two groups.

| Group | KD |  |  |  |  |  |  | Controls |  |
| --- | --- | --- | --- | --- | --- | --- | --- | --- | --- |
| Patients | P1 | P2 | P3 | P4 | P5 | P6 | P7 | C1 | C2 |
| Gender | Male | Male | Male | Female | Female | Female | Male | Male | Female |
| Age (yeras) | 2.1 | 2.0 | 4.7 | 1.6 | 5.3 | 3.3 | 1.9 | 1.1 | 5.3 |
| Duration of fever (days) | 4 | 6 | 4 | 3 | 5 | 6 | 6 | \ | \ |
| Congestion of the bulbar conjunctiva (days) | 1 | 1 | 2 | 3 | 5 | 2 | 5 | \ | \ |
| changes of lips and oral cavity (days) | 1 | 1 | 1 | 3 | 1 | 4 | 5 | \ | \ |
| Rash and erythema multiforme (days) | 2 | 2 | 2 | 3 | 4 | 1 | 6 | \ | \ |
| Swelling of the extremities (days) | 1 | 1 | 1 | 3 | \ | \ | \ | \ | \ |
| Perianal flushing and desquamation (days) | \ | 1 | 1 | 1 | \ | 1 | 1 | \ | \ |
| Enlargement of cervical lymph nodes (days) | 1 | 1 | 1 | 1 | 1 | 1 | 1 | \ | \ |

P: patient, C: control,\:not applicable

Samples collected prior to IVIG treatment were designated as the KDB group. Follow-up blood samples for scRNA-seq analysis were obtained 48 hours after the completion of IVIG therapy, designated as the KDA group. For the Control group, two healthy children (one male and one female, aged between 1.1 and 5.3 years) were enrolled. Detailed clinical information for each participant across the groups is presented in Table 1.

### 3.2 Cellular Composition Analysis of PBMCs in Kawasaki Disease

Single-cell RNA sequencing was performed on the 10x Genomics platform using peripheral blood mononuclear cells isolated from the study participants. After library preparation and stringent quality control, 114,984 high-quality cells were recovered: 103,407 from pre-treatment KD patients and 11,577 from controls. Multidimensional scaling showed distinct clustering by group, indicating clear transcriptomic separation between conditions (supplementary Figure 1).Unsupervised clustering is displayed in two-dimensional space in Figure 1A. Populations were annotated with SingleR and canonical markers. The main PBMC components were CD4+ T cells (28.6 %), CD8+ T cells (12.3 %), other T cells (3.2 %), B cells (19.8 %), NK cells (5.0 %), monocytes (10.0 %), dendritic cells (0.34 %), neutrophils (3.7 %), platelets (2.1 %), erythroblasts (0.3 %), and HSPCs (0.1 %).Neutrophil proportions tended to increase during acute KD (Wilcoxon rank-sum test, P = 0.056), and plasma-cell proportions rose significantly after IVIG treatment (P = 0.036).

**Figure 1:**
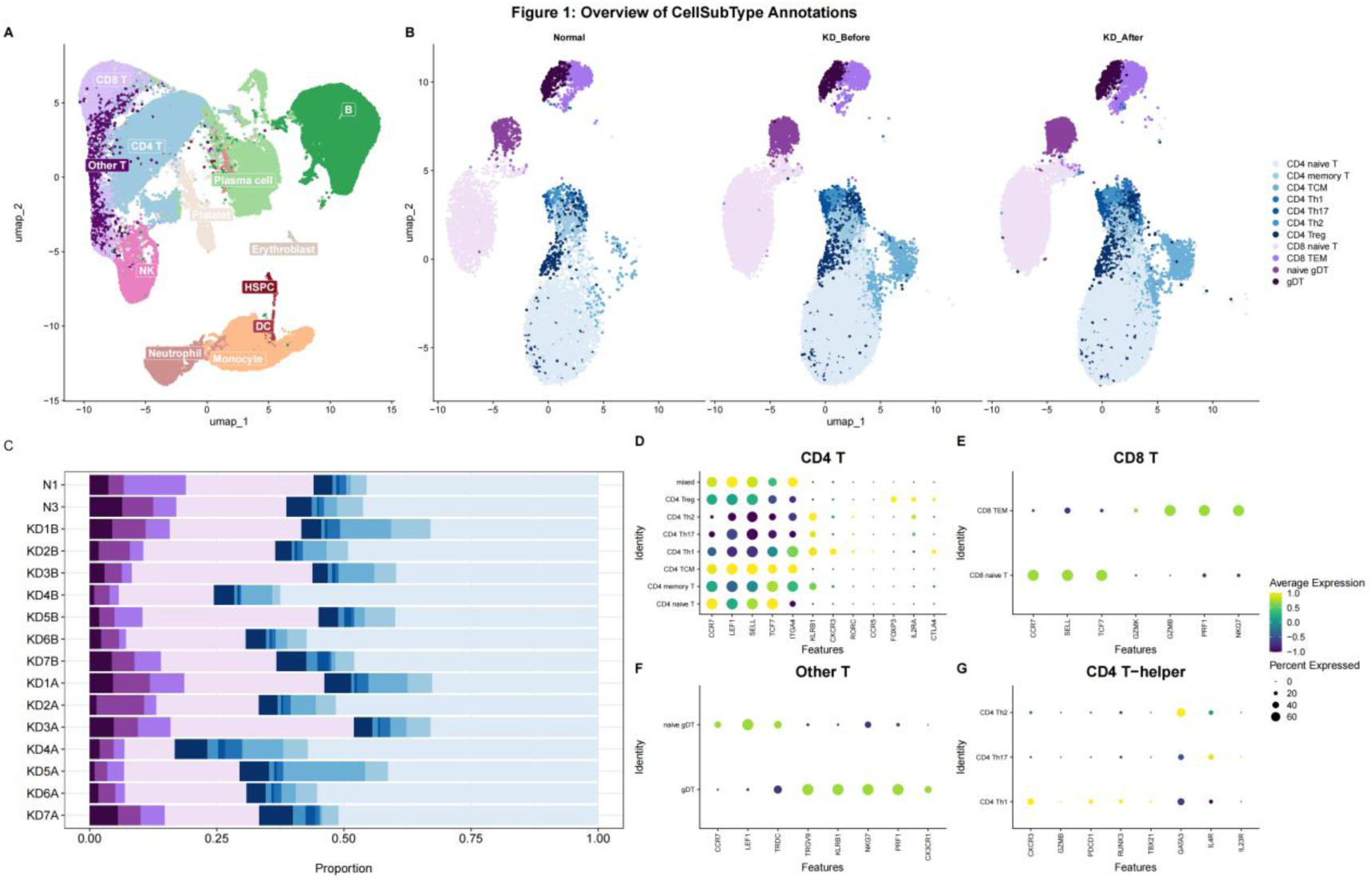
Annotation and Proportional Analysis of Major Immune Cell Clusters and T-cell Subsets. A: Two-dimensional UMAP projection illustrating the major immune cell populations identified across the integrated datasets. B: UMAP visualization showing the distribution of T-cell subsets across three experimental groups: Normal (Healthy Controls), KD_Before (Kawasaki Disease pre-treatment), and KD_After (Kawasaki Disease post-treatment). C: Quantitative compositional analysis of T-cell subsets across individual samples. The stacked bar chart depicts the relative proportions of various T-cell sub-populations in the control group (H1–H2), pre-treatment KD samples (KD1B–KD7B), and post-treatment KD samples (KD1A–KD7A). D–F: Dot plots illustrating the expression of signature marker genes for distinct T-cell subsets. The dot size represents the percentage of cells expressing the gene, and the color intensity indicates the average expression level. D: CD4+ T-cell subsets; E: CD8+ T-cell subsets; F: Other T-cell populations (γδ T-cell lineage); **G:** Heatmap showing Z-score normalized expression of marker genes for CD4+ helper T-cell (Th) subtypes; higher color intensity indicates stronger expression.

### 3.3 Refined Sub-clustering and Annotation of T-cell Subsets

A more detailed sub-clustering analysis was performed to dissect the heterogeneity of T cells, identifying 13 distinct functional subtypes, as illustrated in Figure 1B. These populations included various CD4^+^ T-cell subsets (CD4^+^ Naive T, CD4^+^ Memory T, CD4^+^ TCM, CD4^+^ Th1, CD4^+^ Th17, CD4^+^ Th2, and CD4^+^ Treg), CD8^+^ T-cell subsets (CD8^+^ Naive T and CD8^+^ TEM), and other T-cell lineages (including Naive γδ T and effector γδ T cells). Quantitative compositional analysis of these subtypes across the Normal, KDB, and KDA groups, as well as within individual samples, is illustrated in Figure 1C. The identity of each CD4^+^ T-cell sub-population was validated by the expression of canonical marker genes ^[22]^. Naive CD4^+^ T cells were identified by high expression of CCR7, SELL, TCF7, and LEF1. Central memory CD4^+^ T cells (CD4+ TCM) showed high expression of CCR7, SELL, and ITGA4. Regulatory T cells (Tregs) exhibited characteristic high expression of FOXP3, IL2RA, and CTLA4. Helper CD4^+^ T cells (Th) were defined by the expression of KLRB1, CXCR3, RORC, and CCR5, while KLRB1+ memory CD4+ T cells demonstrated elevated levels of ITGA4 and KLRB1. Through refined resolution of the Th/TEM mixed population, multiple helper T-cell lineages were accurately delineated, including: Th1 cells (CXCR3, GZMB, PDCD1, RUNX3, TBX21), Th2 cells (GATA3+), and Th17 cells (IL23R+). In parallel, independent clustering clarified the major functional states of CD8+ T cells and γδ T cells. The CD8+ T-cell compartment was predominantly composed of Naive CD8^+^ T cells (CCR7+, SELL+, TCF7+) and effector memory CD8^+^ T cells (TEM, GZMK+, GZMB+, PRF1+, NKG7+). Within the CD4^−^CD8^−^ T-cell population, γδ T cells (gDT) were successfully identified and further partitioned into Naive gDT (CCR7+, LEF1+, TRDC+) and Effector gDT (TRGV9+, KLRB1+, NKG7+, PRF1+, CX3CR1+) based on their respective differentiation markers(Figure 1D).

### 3.4 Intercellular Communication Patterns Across Subsets

Systematic analysis of the peripheral immune cell communication landscape across distinct pathological stages revealed that the acute phase of KD is characterized by an exceptionally robust and complex global interaction network (P < 0.05, Permutation test with 100 iterations). The frequency and intensity of communication during this stage were significantly higher than those observed in the control and post-treatment groups(Supplementary Figure S2A, C). This enhanced communication potential was primarily enriched among B cells, monocytes, and effector T cells (Figure 2A). Notably, while B cells exhibited a high predicted signal-radiating capacity, communication effects between certain highly differentiated terminal effector subsets were relatively weak. During the acute phase, the MIF (Macrophage Migration Inhibitory Factor) pathway established an extensive pro-inflammatory interaction network, with B cells predicted as the primary Senders radiating signals toward monocytes and neutrophils (Figure 2B). Comparative analysis of relative overall signaling flow intensity across the three groups is presented in Supplementary Figure S2B and D.Network role analysis (NetAnalysis) indicated that B cells contributed significantly to the outgoing signaling strength, while monocytes acted as the primary receivers, potentially associated with downstream pro-inflammatory responses (Figure 2C).Following IVIG therapy, the overall interaction flux of the MIF pathway underwent a dramatic contraction. Centrality scores for key nodes decreased significantly, and the previously tight “B cell-monocyte” interaction axis was disrupted. These findings suggest that IVIG may facilitate the resolution of systemic inflammation by remodeling MIF-mediated cellular dialogues, thereby steering the immune system toward the restoration of immunological homeostasis.

**Figure 2:**
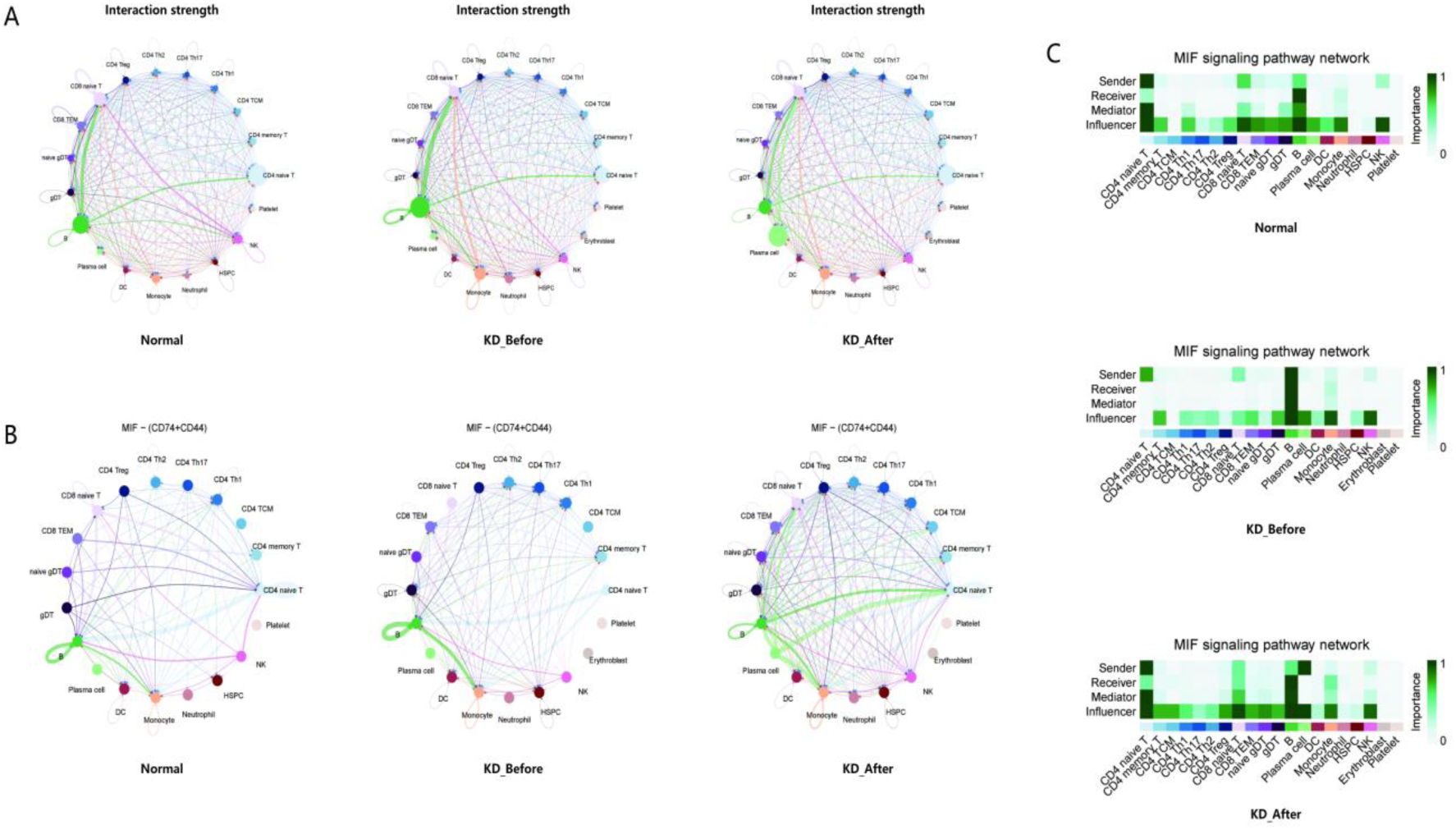
Global Immune Cell Communication Landscape and Dynamic Evolution of the MIF Signaling Pathway during KD Progression. A: Global intercellular communication networks. The hierarchical circle plots illustrate the global crosstalk intensity among peripheral immune cell populations in the Normal Control, KD_Before, and KD_After groups. Circles represent distinct cell populations, while connecting lines indicate interactions; line thickness is proportional to the overall interaction strength.Statistical significance of interaction strength and frequency was determined by permutation test (100 iterations), with P < 0.05 denoted for all visible links. B: Specific interaction network of the MIF signaling pathway (CD74+CD44 receptor complex). The chord diagrams detail the communication topology of the MIF pathway across different disease stages. During the acute phase of KD, a dense pro-inflammatory interactome is observed, characterized by B cells serving as the primary Senders and monocytes as the predominant Receivers. C: Quantitative assessment of cellular roles within the MIF pathway via Network Centrality Analysis (NetAnalysis). The heatmap’s horizontal axis represents cell subtypes, while the vertical axis depicts the four fundamental communication roles: Sender, Receiver, Mediator, and Influencer. The color intensity reflects the relative importance (centrality score) of each cell population within the signaling network.

### 3.5 Developmental Trajectories of T-cell Subsets Across Different Stages of Kawasaki Disease

To further investigate the transcriptional dynamics and state transition relationships of the identified T-cell subsets, particularly CD4^+^ T cells, Monocle3-based pseudotime trajectory analysis was performed on the Normal, KDB, and KDA groups. The cell density distribution of key T-cell subtypes along the pseudotime axis was quantitatively compared across the three conditions (Figure 3A).Several subsets, including CD4^+^ Naive T, CD4^+^ TCM, CD8^+^ Naive T, and Naive γδ T cells, exhibited distribution curves with overlapping or slightly shifted peaks, suggesting that these populations exist at similar transcriptional baselines despite variations in their proportions. Conversely, CD4^+^ Th1 and CD4^+^ Th2 cells clustered at later pseudotime points, consistent with their characteristics as highly polarized effector states. Notably, the CD4^+^ Th17 subset demonstrated the most significant disparity among the groups: Th17 cells in the KDB group were highly concentrated at the latest pseudotime stage, displaying the most complete and distinct peak distribution compared to the Normal and KDA groups. This finding suggests that during the acute phase of KDB, CD4^+^ T cells undergo robust and synchronized functional reprogramming toward a Th17-like inflammatory phenotype.Simultaneously, Naive γδ T and effector γδ T cells showed marked shifts in their transcriptional states along the pseudotime axis, likely reflecting the phenotypic transition occurring during acute inflammation.In the Normal group, the CD4^+^ T-cell trajectory map exhibited a structured profile, establishing a baseline differentiation flux for healthy peripheral CD4^+^ T cells. However, in the KDB group, the CD4^+^ T-cell trajectory underwent substantial alterations. The spatial arrangement of cell clusters differed markedly from the Normal group, reflecting inflammation-driven phenotypic skewing or intensified activation processes (Figures 3B–G). Despite the involvement of various T-cell subsets in KD, the CD4^+^T cell activation path specifically biases toward the Th17-like state during the acute phase, indicating that Th17-mediated inflammatory responses may play a central role in KD pathogenesis. A comparative analysis of the proportions of various sub-populations within the CD4^+^ T-cell compartment revealed a noticeable expansion of Th17 cells during the acute phase of KD (Figure 3H, P = 0.192, d = 0.62), alongside a downward trend in the Treg population (Figure 3I, P = 0.681, d =0.39).While individual proportion comparisons were constrained by the cohort size, a numerical increase in the Th17/Treg ratio was observed during the acute phase(Figure 3J, P = 0.190, d = 0.87), representing a large effect size that signifies a systemic shift in immune balance. This quantitative redistribution, coupled with the phenotypic transitions observed in pseudotime, strongly suggests an inflammatory skewing and an imbalance in CD4+ T-cell homeostasis during acute KD.

**Figure 3:**
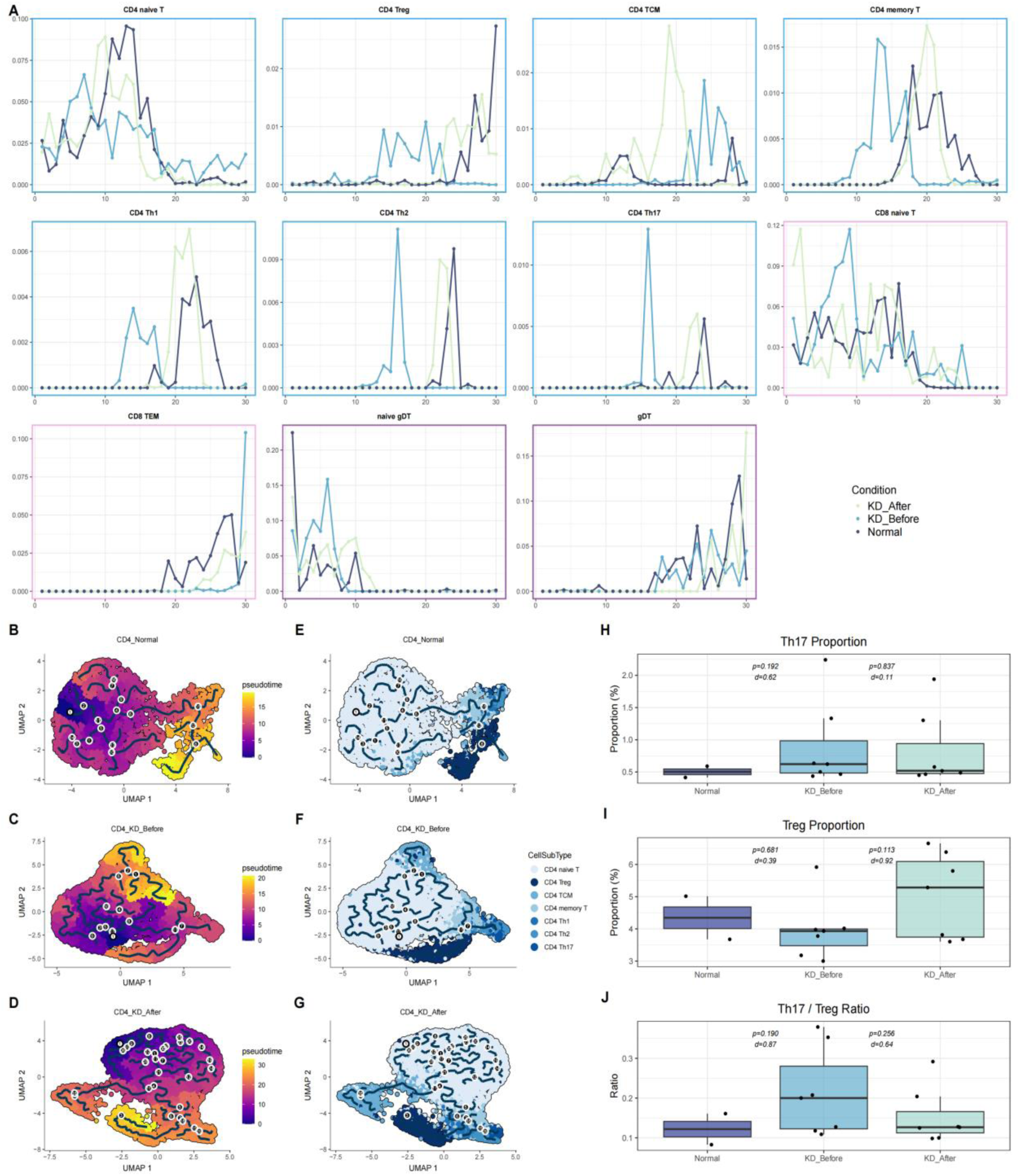
Differentiation Kinetics and Homeostatic Balance Analysis of T-cell Subsets in KD. A: Pseudotime density distribution of key T-cell subtypes across three experimental conditions. The X-axis represents the normalized pseudotime values inferred by Monocle3, and the Y-axis represents cell density. The plots illustrate the distribution fluctuations and enrichment disparities of various T-cell subsets along the developmental trajectory. Note: Y-axis scales for density plots are independently optimized for each CD4 T/CD8 T/Other T cells to normalize for significant differences in abundance among lineages, enabling high-resolution visualization of peak distribution shifts within rare T-cell subtypes along the pseudotime trajectory. B–G: Pseudotime trajectory analysis of CD4+ T cells based on Monocle3. B–D: UMAP projections for the Normal, KD_Before, and KD_After groups, color-coded by pseudotime values. Darker shades indicate earlier developmental stages, while lighter shades represent later stages. E–G: Corresponding cell subtype distribution maps overlaid with the learning graph, revealing the differences in the topological structures of T-cell differentiation paths under different conditions. H–J: Quantitative comparison of T-cell subset proportions and homeostatic balance. H–I: Box plots showing that, compared with the Normal group, the proportion of Th17 cells within the T-cell compartment increased during the acute phase of KD, whereas the proportion of Treg cells exhibited a decreasing trend. J: Analysis of the Th17/Treg ratio, indicating a significant upward trend during the acute phase of KD. * All p-values were calculated using Welch’s t-test to account for unequal variances and sample sizes. Cohen’s d effect sizes are provided to quantify the magnitude of biological differences.

### 3.6 Functional Analysis of T-cell Specialized Expression Modules and TDDGs in Th17-precursor Memory T cells

Pseudotime analysis revealed that acute inflammation in KD triggers a pronounced transcriptional transition in CD4^+^ T cells, characterized by a significant phenotypic shift toward the Th17-like activation state. Consequently, TDDGs identified in CD4^+^ T cells were subjected to functional module analysis. Figure 4A illustrates the overlap of TDDGs among CD4^+^ T cells in the Normal, KDB, and KDA groups. In the KDB group, 1,905 TDDGs (identified by Monocle3 graph_test, q-value < 0.05) were identified (Supplementary Table 1), which are likely associated with the inflammatory phenotype and functional polarization. Based on their co-expression patterns along the CD4^+^ T-cell activation trajectory, these KDB-specific TDDGs were clustered into 17 functional modules (Modules 1–17; Figures 4B, C).

**Figure 4:**
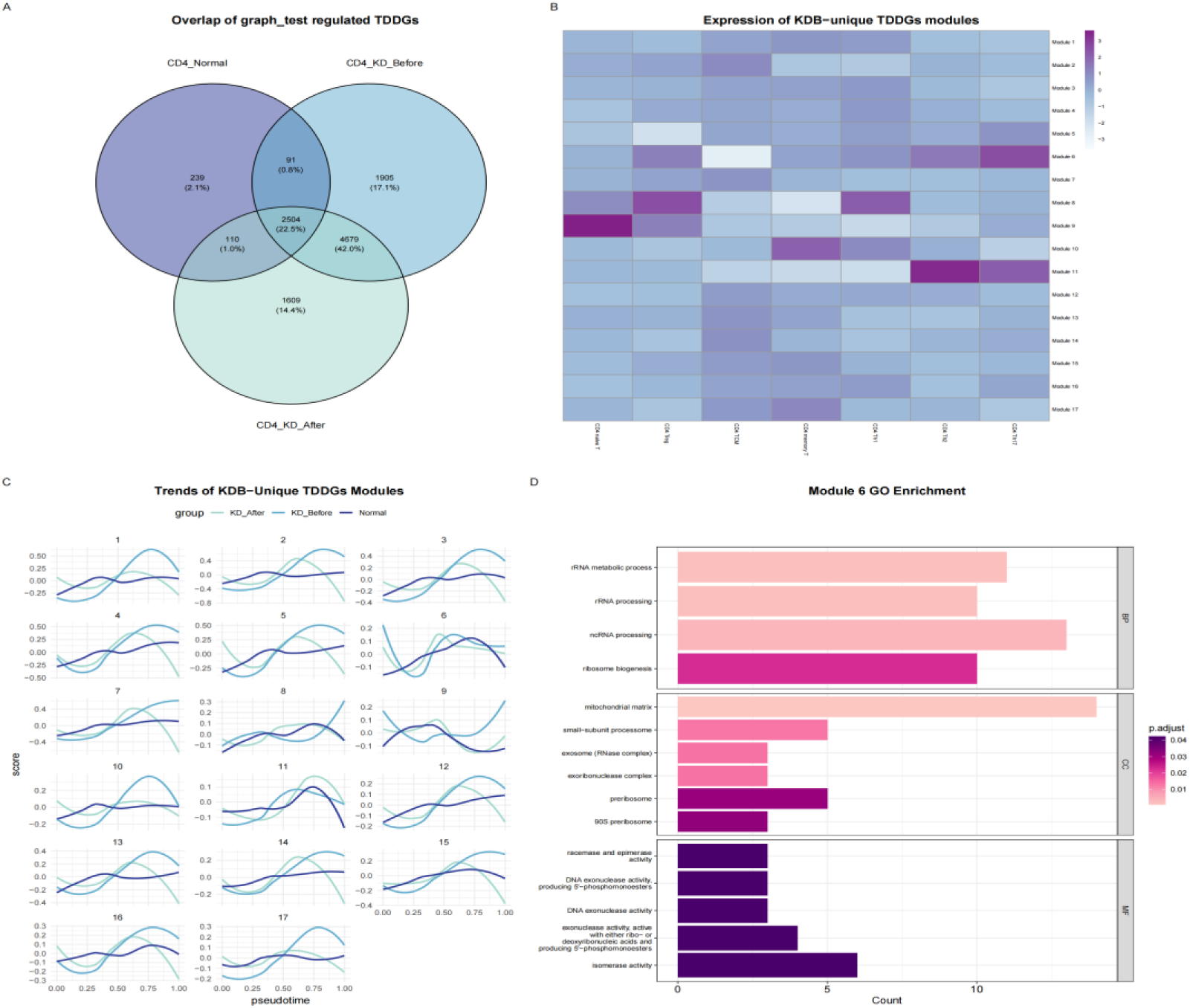
Transcriptional Evolutionary Dynamics of CD4^+^ T-cell Functional Modules during KD Progression. A: Venn diagram analysis of differentially expressed genes across groups. The plot illustrates the numerical distribution and overlap of TDDGs identified in CD4^+^ T cells among the Normal, KD_Before, and KD_After groups.The total number of non-redundant genes identified across three pairwise comparisons (KDB vs. Normal, KDA vs. Normal, and KDB vs. KDA) was used as the universal denominator (Union set, n = 11,141) to calculate the percentage of group-specific or overlapping genes. B: Expression profiles of KD_Before-specific gene clustering modules across T-cell subsets. The heatmap displays the relative expression scores of 17 functional gene modules within each CD4^+^ T-cell sub-population. The color intensity represents the module activity level, revealing the functional heterogeneity of specific cellular phenotypes under acute inflammatory conditions. C: Dynamic expression trends of KD_Before-specific gene modules along the pseudotime trajectory. The line plots demonstrate the aggregate expression kinetics of the 17 functional modules along the normalized pseudotime axis, highlighting their temporal activation patterns during T-cell differentiation.

Among these, Module 6 exhibited a distinct expression profile associated with the activation signature in acute KD: its expression was relatively low in memory T cells but progressively increased across the transition from memory cells to highly activated effector phenotypes, such as Tregs and Th17 cells. Figure 4C clearly shows that Module 6 expression was consistently and significantly upregulated throughout the transcriptional trajectory in the KDB group compared to the Normal group.GO enrichment analysis indicated that the Biological Processes (BP) most significantly enriched in Module 6 were primarily associated with basal cellular biosynthesis and RNA metabolism, reflecting heightened transcriptional and translational activities in activated CD4^+^ T cells. The top-ranked Cellular Components (CC) were closely linked to ribosomal and mitochondrial machinery, while Molecular Functions (MF) emphasized enzymatic activities involved in nucleic acid processing and modification. Functional analysis of Module 6 reveals that the transcriptional shifts in CD4^+^ T cells during acute KD reflect a broad cellular activation signature, characterized by the upregulation of metabolic pathways and ribosomal machinery. Rather than acting as specific drivers of Th17 differentiation, these metabolic and biosynthetic changes likely signify the heightened energetic demands associated with the rapid transition toward an effector phenotype. These findings highlight the metabolic reprogramming that accompanies T-cell functional activation under systemic inflammatory stress.

### 3.7 Transcriptional Signatures Associated with the Th17-like Phenotype

To elucidate the molecular signatures associated with the Th17 phenotypic bias, functional enrichment analysis was performed on the DEGs upregulated in CD4^+^ memory T cells within the KDB group (v.s. Normal group). The results demonstrated that these upregulated genes were significantly enriched in pathways related to T-cell activation, Th17 -related signaling, and metabolic regulation (Figure 5A). Complementary KEGG pathway enrichment analysis further suggested that Th17 cell related processes and the IL-17 effector cytokine production pathway were among the most significantly enriched terms (Figure 5B). These findings indicate that during the acute phase of KD, CD4^+^ memory T cells exhibit a transcriptional profile primed toward Th17-like activation and enhanced metabolic activity. Among the 96 DEGs identified (Supplementary Table 2), key genes associated with the STAT3/JAK signaling axis were upregulated in the acute KD phase, including STAT3 (P < 0.05) and JAK2 (P < 0.05). Similarly, MYC (P < 0.05), associated with cellular metabolic adaptation, showed a significant increase. The expression of RORC (P = 0.41) also exhibited a similar upward trend, although it did not reach statistical significance. Collectively, these findings suggest that the Th17-like transition is associated with the activation of signaling and metabolic programs, supported by the observed expression trends of these key factors. Notably, SOCS3 (P < 0.05), a well-known negative regulator of the JAK/STAT pathway, was also upregulated in the KDB group, likely acting as a compensatory feedback mechanism (Figure 5C). Collectively, these discoveries suggest that the Th17-like transition observed in acute KD is associated with the synchronized transcriptional upregulation of genes within the STAT3/JAK signaling pathway and metabolic programs mediated by factors such as MYC and RORA.

**Figure 5:**
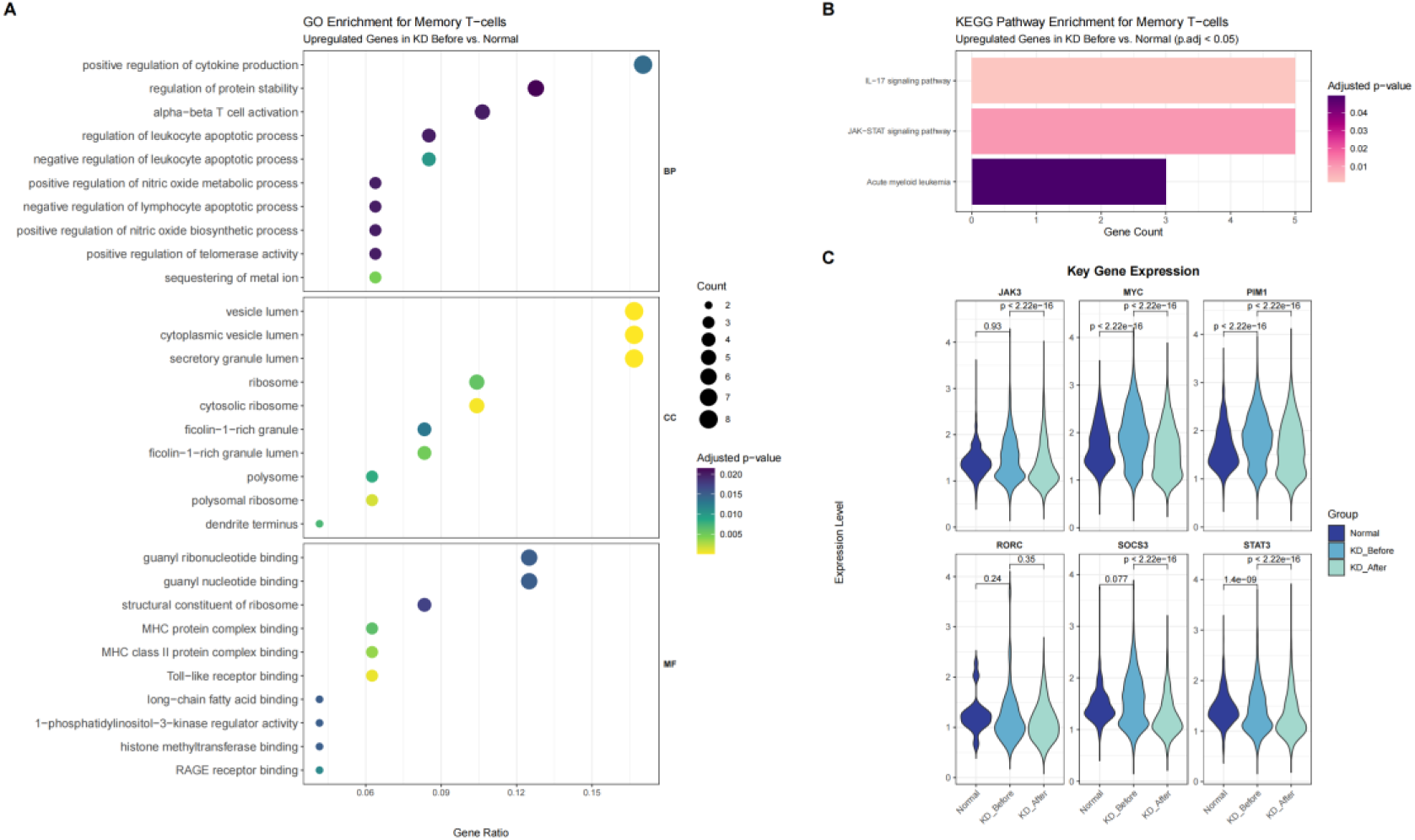
Functional Characteristics and Key Regulatory Gene Analysis of CD4^+^ Memory T Cells During the Acute Phase of KD. A: Gene Ontology enrichment analysis of upregulated differentially expressed genes in CD4^+^ memory T cells. The bubble plot illustrates the significantly enriched terms in Biological Processes, Cellular Components, and Molecular Functions for upregulated genes in the KD_Before group relative to the Normal group (Top 10 terms, adjusted P < 0.05). The bubble size correlates with the gene count, while the color gradient represents the adjusted P-value. B: KEGG pathway enrichment analysis of upregulated DEGs in CD4^+^ memory T cells (adjusted P < 0.05). The horizontal axis represents the gene count enriched within each pathway, and the vertical axis lists the specific KEGG pathway names. The color intensity of the bars maps to the adjusted P-value, with lighter shades indicating higher statistical significance (smaller P-values). C: Expression profiles of key regulatory molecules. Violin plots quantify the expression levels of six core genes across the Normal, KD_Before, and KD_After groups. To ensure clarity, cells with zero expression for the corresponding genes were excluded from the visualization.Statistical significance was determined using the Wilcoxon rank sum test for pairwise comparisons.

**Figure 6:**
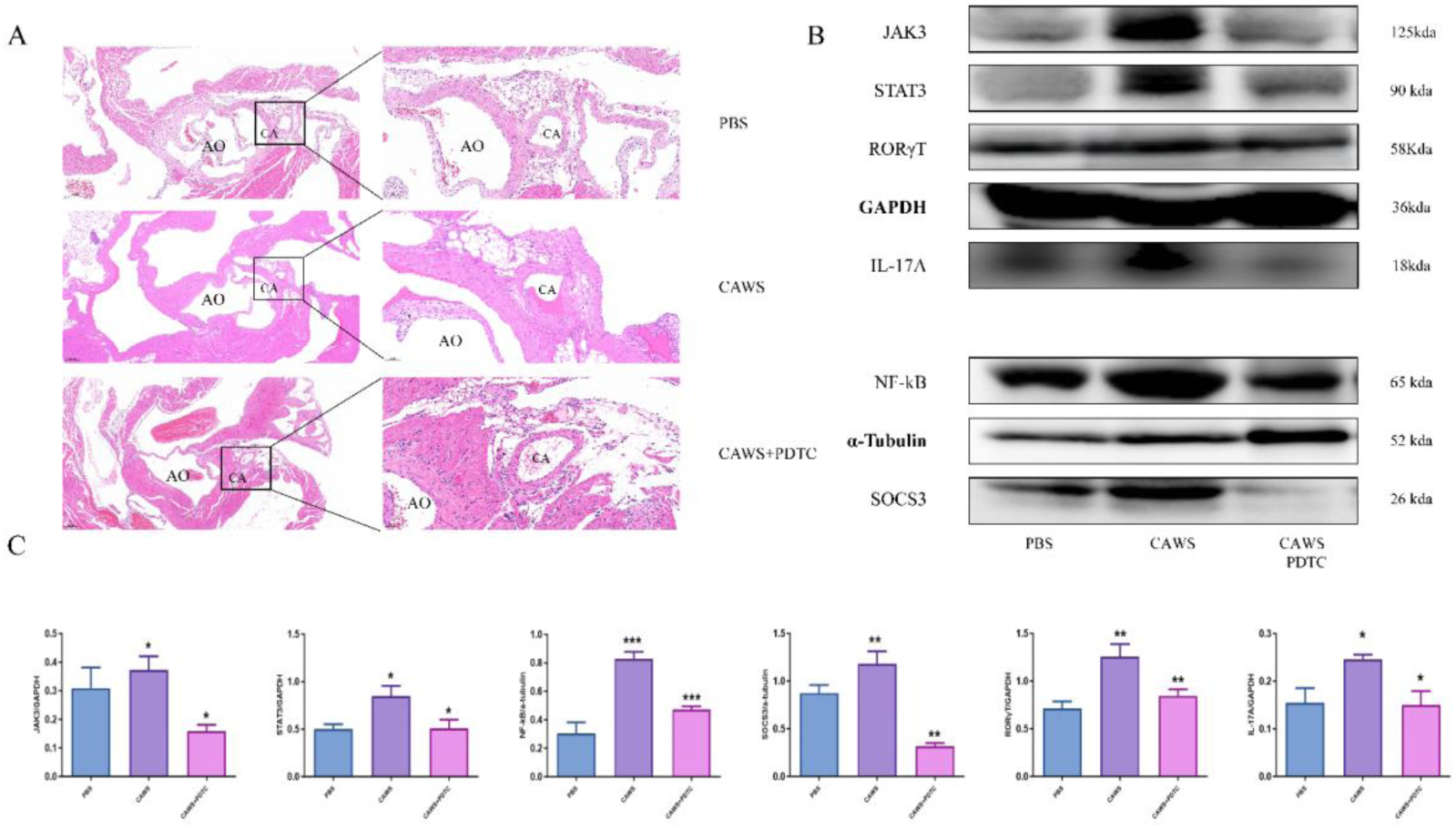
Th17 Cell Inhibitor PDTC Alleviates CAWS-Induced Vasculitis and Blocks the JAK3-STAT3 Signaling Axis in Mice. A: H&E staining of the murine cardiac base and coronary artery tissues. Representative images are shown for the PBS control group, CAWS model group, and CAWS+PDTC treatment group. AO denotes the aortic root, and CA denotes the coronary artery. The images on the right provide high-magnification views of the corresponding regions. B: Western blot analysis was used to detect changes in the STAT3/JAK3 signaling pathway and the protein expression levels of inflammatory factors NF-κB and IL-17a in mice across different groups. C: Quantitative statistical analysis of protein expression. The gray values were quantified using ImageJ. Data are presented as mean ± SEM in bar graphs. Note: n = 6; *: P < 0.05; **: P < 0.01; ***: P < 0.001.

### 3.8 Establishment of the KD Animal Model and Attenuation of Th17-associated Responses on Vascular Injury

The histopathological hallmarks of vascular lesions in the CAWS mouse model mirrored those observed in human KD, with inflammation concurrently affecting the aortic root and the perivascular regions of the coronary arteries. Hematoxylin and eosin staining revealed that at 28 days, the CAWS group exhibited significant inflammatory cell infiltration within the vessel walls of the aortic root and coronary arteries, showing a distribution highly consistent with the histological changes in KD patients. Following pharmacological intervention with PDTC, a potent anti-inflammatory and antioxidant agent, inflammatory infiltration in the corresponding areas was markedly reduced, whereas the PBS group maintained intact vascular architecture with virtually no inflammatory response.

The effects of PDTC intervention on inflammatory signaling were further investigated in the CAWS vasculitis model. Western blot analysis demonstrated that, compared to the PBS group, the expression levels of pathway proteins—including STAT3, JAK3, SOCS3, and RORγt—as well as inflammatory markers IL-17A and NF-κB, were significantly elevated in the cardiac tissues of the CAWS group. This indicates heightened inflammatory signaling is a prominent feature in the CAWS mouse model. Conversely, compared to the CAWS group, the CAWS+PDTC group exhibited a significant downregulation in the expression of STAT3, JAK3, SOCS3, RORγt, IL-17A, and NF-κB, suggesting that PDTC-mediated anti-inflammatory effects are associated with the suppression of multiple signaling axes, including STAT3/JAK3 and NF-κB.Overall, the results demonstrate that mitigating the systemic inflammatory response can effectively reduce the expression of these core proteins, thereby alleviating inflammatory activation and vascular damage in CAWS mice.

## 4. Discussion

This study provides a high-resolution annotation of T lymphocytes in the peripheral blood of patients with KD using single-cell RNA sequencing (scRNA-seq). Given the high heterogeneity of T cells, a strategy of hierarchical clustering and targeted annotation was employed to construct the most detailed T-cell immune landscape associated with KD to date. Deep analysis revealed that CD4^+^ T cells in the acute phase are highly enriched with specific transcriptional programs related to the Th17 lineage. Previous studies have indicated that the Th17/Treg balance is disrupted in KD patients and is closely associated with resistance to intravenous immunoglobulin (IVIG) therapy ^[5, 23]^. The current findings demonstrate a significant transcriptional shift toward a Th17-like state in acute-phase CD4+ T cells, a trend consistent with elevated systemic inflammatory cytokines and the enhancement of the downstream IL-17 signaling pathway.

The core effector cytokine secreted by Th17 cells, IL-17A, is a key mediator of vascular inflammation in KD ^[24]^. Under pathological conditions, IL-17A acts directly on vascular endothelial cells (VECs) to induce the expression of adhesion molecules such as ICAM-1 and VCAM-1 by activating downstream pathways such as NF-κB or MAPK, thereby promoting the recruitment and infiltration of neutrophils and monocytes into the vascular wall ^[25, 26]^. Furthermore, IL-17A induces the secretion of various chemokines (e.g., CXCL1, CXCL8) by endothelial cells and acts synergistically with TNF-α to amplify endothelial inflammatory feedback, ultimately leading to the upregulation of matrix metalloproteinases (MMPs) and degradation of the vascular basement membrane ^[25, 27, 28]^. Significant enrichment of IL-17 signaling pathway genes during the acute phase suggests that the activation of this axis may play a crucial role in the early initiation of coronary artery damage in KD.

The study delineates a temporal progression from an early myeloid-driven microenvironment to late-stage T-cell effector differentiation in KD. Cell-cell communication analysis highlights the involvement of the MIF axis in the formation of the initial inflammatory milieu, while pseudotime analysis reveals that CD4+ T cells subsequently undergo Th17 polarization via the JAK/STAT3 pathway. This logical transition from early extrinsic priming to intrinsic transcriptional reprogramming underscores the orchestration of systemic inflammation and localized immune damage in KD.

STAT3 serves as a central molecular switch regulating Th17-like functions ^[29]^. In this study, significantly elevated expression of STAT3, JAK2, and SOCS3 was observed in acute-phase memory T cells. This STAT3-centered signaling cascade not only drives the directional differentiation of pro-inflammatory phenotypes but also profoundly reshapes cellular metabolic programs through direct transcriptional regulation. Previous research has confirmed that phosphorylated STAT3 can bind directly to the promoter regions of rate-limiting glycolytic enzymes, such as hexokinase 2 (HK2) and pyruvate kinase M2 (PKM2), inducing their high expression to initiate aerobic glycolysis ^[30–32]^. Concurrently, synchronized enhancement in the expression of MYC, a key regulator of glycolysis, was observed. Under inflammatory stress, STAT3 can synergistically induce MYC transcription by recruiting co-activators. Subsequently, MYC upregulates the expression of glucose transporters (e.g., GLUT1), forming a cascade effect in energy metabolism ^[33] [34]^. This STAT3-driven metabolic reprogramming provides the necessary material basis and energy support for T cells to maintain explosive proliferative capacity and massive secretion of IL-17A in an inflammatory environment ^[35]^. Therefore, STAT3 likely mediates not only cell differentiation but also provides metabolic support for the continuous amplification of the inflammatory response by regulating T-cell metabolic reprogramming during the acute phase of KD. Reflecting the cascade effects of the STAT3 pathway, intervention targeting the JAK/STAT3 axis has become a focal point of clinical research. Studies have shown that JAK inhibitors exhibit promising potential in patients with juvenile idiopathic arthritis (JIA) ^[36]^. Their efficacy may stem from the broad-spectrum blockade of upstream signals, thereby inhibiting STAT3-induced Th17 differentiation and metabolic activation ^[37]^. Additionally, targeted IL-17A inhibitors, such as Secukinumab, have demonstrated potential in protecting endothelial function in systemic vasculitis ^[38]^. In the CAWS mouse model, targeted blockade of STAT3 and its downstream IL-17A expression by PDTC resulted in a significant reduction of vascular injury. These results collectively suggest that precise intervention targeting the STAT3 cascade and its downstream IL-17 axis represents a potential innovative direction for alleviating vascular inflammation and protecting the coronary endothelium in KD.

In conclusion, this study characterizes the transcriptional landscape of peripheral T cells in acute KD and highlights the potential role of STAT3-driven metabolic reprogramming and Th17 differentiation in mediating endothelial injury, providing a theoretical basis for understanding KD pathogenesis and developing novel therapeutic strategies.

Several limitations remain in this study. Pseudo-time analysis reflects cell state trends rather than confirmed differentiation trajectories. Inferences based on peripheral blood transcriptomes are speculative, and peripheral blood data may not fully represent the local microenvironment of coronary artery infiltration. Future validation using spatial transcriptomics and in-depth in vitro and in vivo mechanistic experiments is required.

---

**Supplementary Figure 1:**
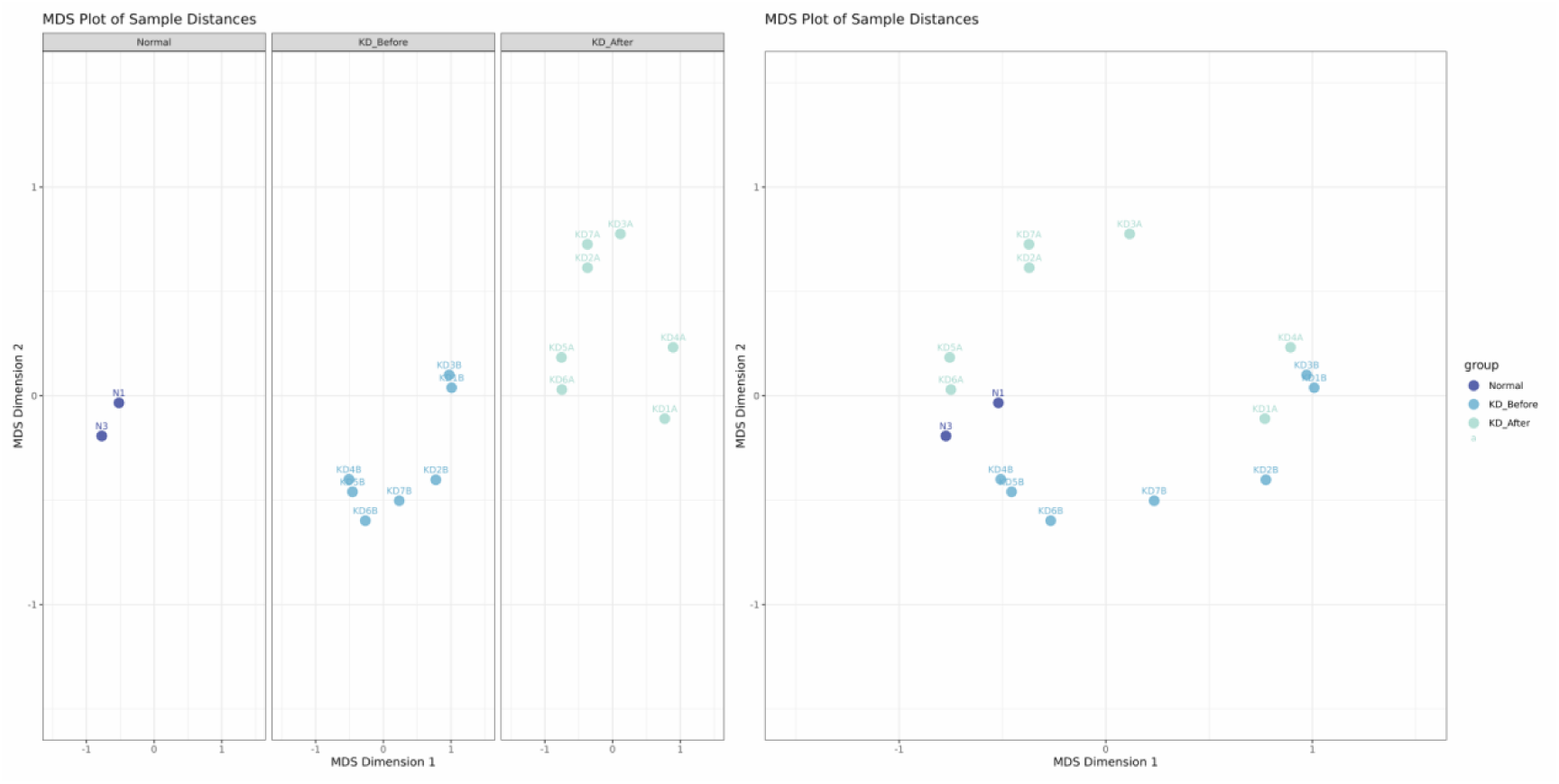
Multidimensional Scaling (MDS) Scatter Plot Based on Transcriptomic Data. The horizontal and vertical axes represent the two primary dimensions (MDS Dimension 1 & 2) that capture the greatest transcriptomic variance among samples. Each data point is color-coded to distinguish between the three experimental cohorts: Normal (Healthy Controls), KD_Before (Acute Kawasaki Disease), and KD_After (Post-treatment).

**Supplementary Figure 2:**
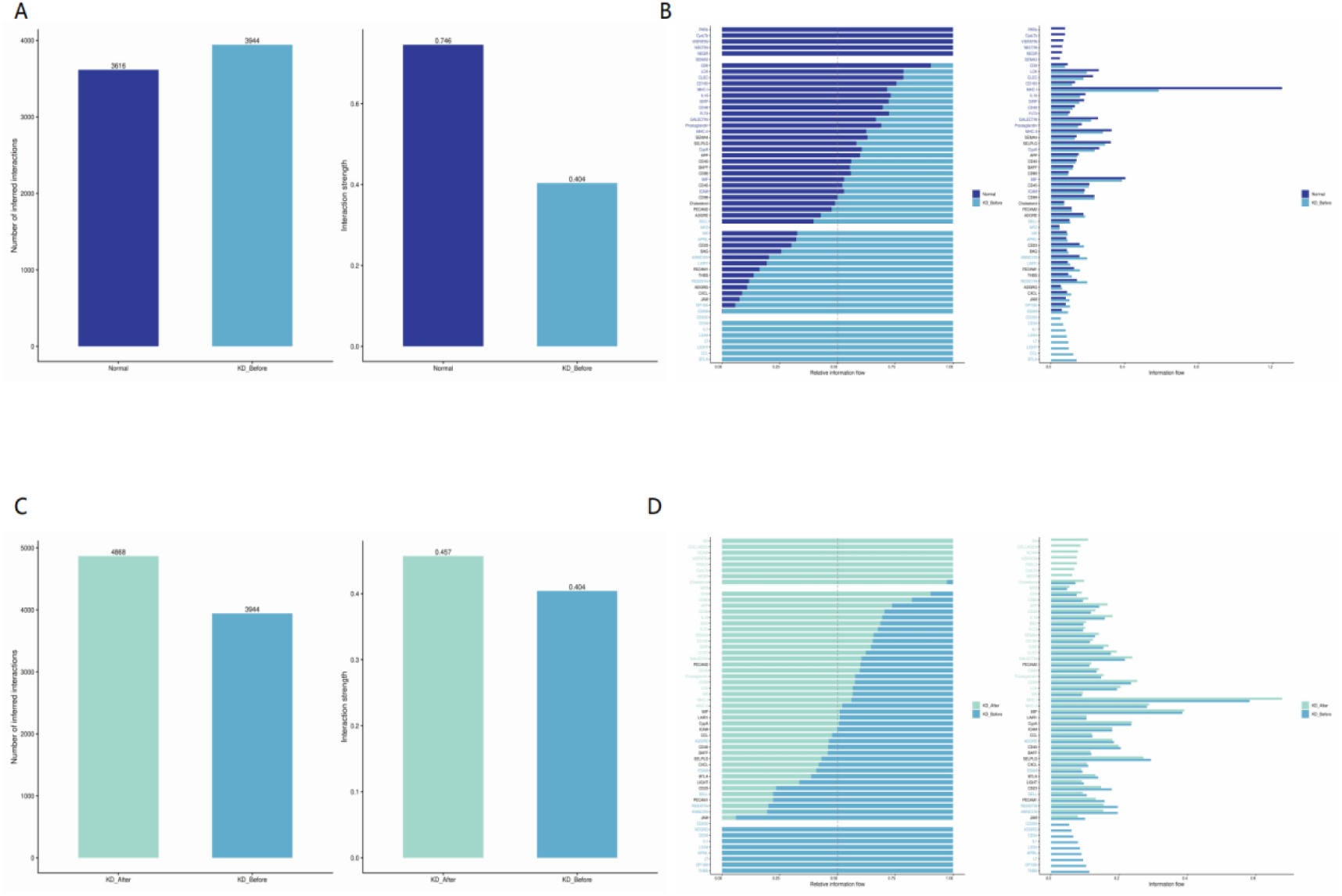
Quantitative comparison and pathway ranking of global cell-cell communications between different groups. (A, C) Interaction frequency and aggregated weight; (B, D) Information flow ranking of significant signaling pathways. *Comparisons: (A, B) KD-Before vs. Normal. (C, D) KD-Before vs. KD-After.

## Supplemental Table legends

**Supplemental table 1.** Identification of Specifically Regulated Unique Genes in the KDB Group.

**Supplemental table 2.** Upregulated DEGs and GO enrichment in CD4+ memory T cells (KD-Before vs. Normal).

**Supplemental table 3.** Donor-level DESeq1 statiscal validation of target genes bewteen Normal group and KD-Before group.

**Supplemental table 4.** Donor-level DESeq1 statiscal validation of target genes bewteen KD-After group and KD-Before group.

**Supplemental table 5.** Expression statistics of lineage marker genes across T-cell subtypes.

## Acknowledgements

We wish to thank all the donors who contributed samples.We thank Dr. Yu-e Guo of the Department of Pharmacy, Shanghai Pulmonary Hospital, Tongji University School of Medicine, Shanghai, China, for generously sharing the PDTC-based Th17-blocking protocol that underpinned this study.

## Author contributions

SS presented the main idea and wrote the manuscript. LqC and YyZ collected the clinical samples. SS, YX analyzed the data. YX and GL revised the manuscript. TX and MH provided the guidance. YfZ,SS,LC completed the basic experimental work.All authors provided intellectual input and read the manuscript.

## Source of Funding Support

National Natural Science Foundation of China (82370296)

Shanghai Municipal Science and Technology Commission Science and Technology Innovation Action Plan’ Medical Innovation Research Special Project (21Y31900304)

Shanghai Municipal Health Commission Medical New Technology Research and Transformation Seed Program (2024ZZ1024).

## Declarations

The study has been reviewed and approved by the Ethics Committee of Shanghai Children’s Hospital (No.2019R081E01).

Trial registration: ChiCTR, ChiCTR2100044729 Registered 24 December 2019, http://www.medresman.org.cn/pub/cn/proj/projectshow.aspx?proj=7739

## Consent for publication

Written informed consent for publication was obtained from all participants prior to the enrollment of this study.

## Competing interests

No potential conflict of interest was reported by the authors

## Data availability

Raw and processed sequence data from newly included samples P1, have been deposited in the National Omics Data Encyclopedia database of Bio-Med Big Data Center, Shanghai Institute of Nutrition and Health, Chinese Academy of Sciences under accession code OEP005481.P2,P3,P4,P5,P6,P7,H1,H2 derived from the samples deposited previously. The raw data generated have been deposited in the National Omics Data Encyclopedia database of Bio-Med Big Data Center, Shanghai Institute of Nutrition and Health, Chinese Academy of Sciences under accession code OEP001162.The processed sequencing data generated have been deposited in the Gene Expression Omnibus (GEO) database under accession code GSE1687320.The raw sequence data are available under restricted access because of data privacy laws, and access can be obtained by reasonable request to the corresponding authors.

## Notes

### Competing Interest Statement

The authors have declared no competing interest.

## References

[1] Jone PN, Tremoulet A, Choueiter N, et al. Update on Diagnosis and Management of Kawasaki Disease: A Scientific Statement From the American Heart Association. Circulation. 2024. 150(23): e481–e500.

[2] Takahashi K, Oharaseki T, Naoe S, Wakayama M, Yokouchi Y. Neutrophilic involvement in the damage to coronary arteries in acute stage of Kawasaki disease. Pediatr Int. 2005. 47(3): 305–10.

[3] Makino N, Nakamura Y, Yashiro M, et al. Nationwide epidemiologic survey of Kawasaki disease in Japan, 2015-2016. Pediatr Int. 2019. 61(4): 397–403.

[4] Lee SB, Kim YH, Hyun MC, Kim YH, Kim HS, Lee YH. T-Helper Cytokine Profiles in Patients with Kawasaki Disease. Korean Circ J. 2015. 45(6): 516–21.

[5] Jia S, Li C, Wang G, Yang J, Zu Y. The T helper type 17/regulatory T cell imbalance in patients with acute Kawasaki disease. Clin Exp Immunol. 2010. 162(1): 131–7.

[6] Noval Rivas M, Lee Y, Wakita D, et al. CD8+ T Cells Contribute to the Development of Coronary Arteritis in the Lactobacillus casei Cell Wall Extract-Induced Murine Model of Kawasaki Disease. Arthritis Rheumatol. 2017. 69(2): 410–421.

[7] Wang S, Qian G, Liu Y, et al. Kawasaki disease: insights into the roles of T cells. Front Immunol. 2025. 16: 1582638.

[8] Wang Z, Xie L, Ding G, et al. Single-cell RNA sequencing of peripheral blood mononuclear cells from acute Kawasaki disease patients. Nat Commun. 2021. 12(1): 5444.

[9] Song S, Chen L, Zhou Y, et al. CD14(+) monocytes: the immune communication hub in early vasculitis symptoms of Kawasaki disease. Front Immunol. 2025. 16: 1557231.

[10] Song S, Chen L, Zhou Y, et al. Single cell sequencing technology reveals the correlation between B lymphocytes and vascular inflammatory symptoms of Kawasaki disease. Transl Pediatr. 2025. 14(4): 582–596.

[11] McCrindle BW, Rowley AH, Newburger JW, et al. Diagnosis, Treatment, and Long-Term Management of Kawasaki Disease: A Scientific Statement for Health Professionals From the American Heart Association. Circulation. 2017. 135(17): e927–e999.

[12] Amezquita RA, Lun A, Becht E, et al. Orchestrating single-cell analysis with Bioconductor. Nat Methods. 2020. 17(2): 137–145.

[13] Hao Y, Stuart T, Kowalski MH, et al. Dictionary learning for integrative, multimodal and scalable single-cell analysis. Nat Biotechnol. 2024. 42(2): 293–304.

[14] McGinnis CS, Murrow LM, Gartner ZJ. DoubletFinder: Doublet Detection in Single-Cell RNA Sequencing Data Using Artificial Nearest Neighbors. Cell Syst. 2019. 8(4): 329–337.e4.

[15] Korsunsky I, Millard N, Fan J, et al. Fast, sensitive and accurate integration of single-cell data with Harmony. Nat Methods. 2019. 16(12): 1289–1296.

[16] Zappia L, Oshlack A. Clustering trees: a visualization for evaluating clusterings at multiple resolutions. Gigascience. 2018. 7(7): giy083.

[17] Jin S, Guerrero-Juarez CF, Zhang L, et al. Inference and analysis of cell-cell communication using CellChat. Nat Commun. 2021. 12(1): 1088.

[18] Cao J, Spielmann M, Qiu X, et al. The single-cell transcriptional landscape of mammalian organogenesis. Nature. 2019. 566(7745): 496–502.

[19] Xu S, Hu E, Cai Y, et al. Using clusterProfiler to characterize multiomics data. Nat Protoc. 2024. 19(11): 3292–3320.

[20] Nagi-Miura N, Harada T, Shinohara H, et al. Lethal and severe coronary arteritis in DBA/2 mice induced by fungal pathogen, CAWS, Candida albicans water-soluble fraction. Atherosclerosis. 2006. 186(2): 310–20.

[21] Guo YE, Lv J, Shu P, et al. Drug screening identifies pyrrolidinedithiocarbamate ammonium ameliorating DSS-induced mouse ulcerative colitis via suppressing Th17 differentiation. Cell Immunol. 2024. 405-406: 104887.

[22] Martin OP, Wallace MS, Oetheimer C, et al. Single-cell atlas of human liver and blood immune cells across fatty liver disease stages reveals distinct signatures linked to liver dysfunction and fibrogenesis. Nat Immunol. 2025. 26(9): 1596–1611.

[23] Hirabayashi Y, Takahashi Y, Xu Y, et al. Lack of CD4⁺CD25⁺FOXP3⁺ regulatory T cells is associated with resistance to intravenous immunoglobulin therapy in patients with Kawasaki disease. Eur J Pediatr. 2013. 172(6): 833–7.

[24] Li H, Xing H. Interleukin-35 Enhances Regulatory T Cell Function by Potentially Suppressing Their Transdifferentiation into a T Helper 17-Like Phenotype in Kawasaki Disease. Immunol Invest. 2023. 52(4): 513–528.

[25] Robert M, Miossec P, Hot A. The Th17 Pathway in Vascular Inflammation: Culprit or Consort. Front Immunol. 2022. 13: 888763.

[26] Nguyen H, Chiasson VL, Chatterjee P, Kopriva SE, Young KJ, Mitchell BM. Interleukin-17 causes Rho-kinase-mediated endothelial dysfunction and hypertension. Cardiovasc Res. 2013. 97(4): 696–704.

[27] Zhang Z, Zhao L, Zhou X, Meng X, Zhou X. Role of inflammation, immunity, and oxidative stress in hypertension: New insights and potential therapeutic targets. Front Immunol. 2022. 13: 1098725.

[28] Li G, Zhang Y, Qian Y, et al. Interleukin-17A promotes rheumatoid arthritis synoviocytes migration and invasion under hypoxia by increasing MMP2 and MMP9 expression through NF-κB/HIF-1α pathway. Mol Immunol. 2013. 53(3): 227–36.

[29] Poholek CH, Raphael I, Wu D, et al. Noncanonical STAT3 activity sustains pathogenic Th17 proliferation and cytokine response to antigen. J Exp Med. 2020. 217(10): e20191761.

[30] Shi LZ, Wang R, Huang G, et al. HIF1alpha-dependent glycolytic pathway orchestrates a metabolic checkpoint for the differentiation of TH17 and Treg cells. J Exp Med. 2011. 208(7): 1367–76.

[31] Damasceno L, Prado DS, Veras FP, et al. PKM2 promotes Th17 cell differentiation and autoimmune inflammation by fine-tuning STAT3 activation. J Exp Med. 2020. 217(10): e20190613.

[32] Demaria M, Giorgi C, Lebiedzinska M, et al. A STAT3-mediated metabolic switch is involved in tumour transformation and STAT3 addiction. Aging (Albany NY). 2010. 2(11): 823–42.

[33] Kunkl M, Sambucci M, Ruggieri S, et al. CD28 Autonomous Signaling Up-Regulates C-Myc Expression and Promotes Glycolysis Enabling Inflammatory T Cell Responses in Multiple Sclerosis. Cells. 2019. 8(6): 575.

[34] Barré B, Vigneron A, Coqueret O. The STAT3 transcription factor is a target for the Myc and riboblastoma proteins on the Cdc25A promoter. J Biol Chem. 2005. 280(16): 15673–81.

[35] Ma S, Ming Y, Wu J, Cui G. Cellular metabolism regulates the differentiation and function of T-cell subsets. Cell Mol Immunol. 2024. 21(5): 419–435.

[36] Ruperto N, Brunner HI, Synoverska O, et al. Tofacitinib in juvenile idiopathic arthritis: a double-blind, placebo-controlled, withdrawal phase 3 randomised trial. Lancet. 2021. 398(10315): 1984–1996.

[37] Chikhoune L, Poggi C, Moreau J, et al. JAK inhibitors (JAKi): Mechanisms of action and perspectives in systemic and autoimmune diseases. Rev Med Interne. 2025. 46(2): 89–106.

[38] Venhoff N, Schmidt WA, Bergner R, et al. Safety and efficacy of secukinumab in patients with giant cell arteritis (TitAIN): a randomised, double-blind, placebo-controlled, phase 2 trial. Lancet Rheumatol. 2023. 5(6): e341–e350.

